# TiSAn: Estimating Tissue Specific Effects of Coding and Noncoding Variants

**DOI:** 10.1101/141408

**Authors:** Kévin Vervier, Jacob J. Michaelson

**Author notes:** Corresponding Author: Jacob J Michaelson, 501 Newton Road, Iowa City, IA 52242.319-335-8066 Contact.

## Abstract

Measures of general deleteriousness, like CADD or PolyPhen, have become indispensable tools in the interpretation of genetic variants. However, these measures say little about where in the organism these deleterious effects will be most apparent. An additional, complementary measure is needed to link deleterious variants (as determined by e.g., CADD) to tissues in which their effect will be most meaningful. Here, we introduce TiSAn (Tissue Specific Annotation), a tool that predicts how related a genomic position is to a given tissue (http://github.com/kevinVervier/TiSAn). TiSAn uses machine learning on genome-scale, tissue-specific data to discriminate variants relevant to a tissue from those having no bearing on the development or function of that tissue. Predictions are then made genome-wide, and these scores can then be used to contextualize and filter variants of interest in whole genome sequencing or genome wide association studies (GWAS). We demonstrate the accuracy and versatility of TiSAn by introducing predictive models for human heart and human brain, and detecting tissue-relevant variations in large cohorts for autism spectrum disorder (TiSAn-brain) and coronary artery disease (TiSAn-heart). We find that TiSAn is better able to prioritize genetic variants according to their tissue-specific action than the current state of the art method, GenoSkyLine.

## Introduction

Whole genome sequencing (WGS) is assuming its role as the technology of choice for an increasing number of genetic studies. A vast majority of the information yielded by WGS resides in non-coding and less well-characterized regions of the genome. Recent work in the annotation of non-coding variation has shown that multiple levels of information, integrated using machine learning algorithms, are required to capture the diverse regulatory potentials in these regions(1-4). However, current state-of-the-art variant annotation methods predict generic pathogenicity, and largely avoid the question of which tissues, organs, and systems are likely to be most susceptible to a particular genetic variation. Projects such as the Genotype-Tissue Expression (GTEx)(5) repository and the NIH Roadmap Epigenomics Mapping Consortium (RME)(6), provide clear evidence that a variant will not necessarily have the same impact on gene expression in different tissues or cell types. Recently proposed approaches, such as GenoSkyline(7), have employed cross-tissue methylation levels to annotate genetic variations. However, such methods have limitations because they were trained only using data that was uniformly collected across a wide variety of tissues, leaving out potentially informative features derived from one-off databases for specific tissues. This results in emphasizing performance over many tissues, rather than optimizing for a specific tissue.

In this work, we introduce Tissue Specific Annotation (TiSAn), which combines the power of supervised machine learning, with tissue-specific annotations, including genomics, transcriptomics and epigenomics (Supplementary Software 1 and http://github.com/kevinVervier/TiSAn for the latest version). We describe a general statistical learning framework in which researchers can derive a nucleotide-resolution score for the tissues they focus on. As a proof of principle, we apply our methodology to two human tissues, namely brain and heart, and we have made pre-computed scores for these two tissues available.

## Materials & Methods

### Training set definition

We identify multiple sources for training examples with respect to a given tissue *T.* One way to build such a training set is to look for positions known to be causal of a disease in tissue *T.* This way, we reduce the risk of training a pathogenicity score and make sure we extract a signal orthogonal to deleteriousness. For deriving a genome-wide predictor, the training set needs to cover both coding and non-coding loci, but also loci related to *T* (positive examples) and unrelated (negative examples). A complete list of the positions used for training brain and heart models is provided in Github vignettes (http://github.com/kevinVervier/TiSAn/tree/master/vignettes). Two types of public databases were used to derive training sets:

### Genotype array loci

Disease-related loci could be found in Consortium developed arrays, designed for targeting specific disorders, such as the MetaboChip(8) for cardiovascular diseases, or Illumina Infinium PsychArray Beadchip for psychiatric disorders. These probe sets contain tissue-related variants (positive examples), but also backbone/non-psychiatric variants which we consider as negative examples if they meet a minimal CADD threshold described below.

### Large intergenic non-coding RNAs

Usually, non-coding variants are less functionally characterized than coding ones. Large intergenic non-coding RNAs (lincRNAs) represent a well-study group of non-coding elements. Databases, such as LincSNP(9), contain disease-related variants that occur in lincRNA loci. After defining a list of tissue-related disorders, we propose to divide this database in two subsets: one related to tissue T (positive examples) and one containing background variants (negative examples), i.e., deleterious randomly sampled variants not related to the tissue at hand. This way, we enrich the training set with non-coding loci.

Exact number of training examples used for TiSAn brain and heart models can be found in Supplementary Table 1.

### Weibull distribution

Following Cherkasov et al. (10), we model the distance between a given locus *x* and a known annotation, following the Weibull distribution and its Extreme Value Theory application. Therefore, in the following paragraphs, the distance is measured as:

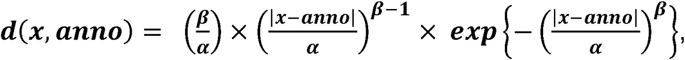

where anno refers to a known annotation position, α is a scale factor, and β is a shape parameter. Parameters fitting was performed separately for each annotation, using *MASS* R package.

### Features extraction

We represent each genomic position in a functional space made of hundreds of different annotations. In the following, we describe how such signal can be extracted using publically available data sets. More details can be found in Github vignettes (http://github.com/kevinVervier/TiSAn/tree/master/vignettes).

Nucleotide frequencies are linked to overall regulatory activity (G/C content), and patterns in nucleotide k-mers are at the core of transcription factor binding site detection(11). Recently, specific patterns have been identified to be tissue-specific(12), and we incorporate this information by computing frequencies for all n-nucleotides (***n*** ∈ (***l,2,3,4***)), found in a +/- 500 base pair neighborhood around a locus ***x***.

Links between disease traits and tissue-specific gene expression have been reported in studies using the rich GTEx dataset(13). For each genomic location, we extract features based on how close X is to known eQTLs for tissue *T*, and for other tissues. Weibull distribution was used for modeling the minimal distance to a GTEx eQTL (Supplementary Fig. 1). We also derive Boolean features for whether or not the genomic position ***x*** is at the exact location of a GTEx eQTL, which puts more weight for being a known locus. Although some genes expression shows variation across tissues, comprehensive resources and exhaustive list of tissue-specific genes are limited. It has already been shown that text mining techniques may help to extract relationship between genes and disease traits(14). Therefore, we propose to adapt such methods to identify, in the Pubmed database (May 2016 gene2ID database), genes reported to be associated with tissue *T.* Only genes with at least 3 citations were kept. We observed, when training the brain model, that around 1,000 tissue-related genes represent enough genome coverage to derive a feature based on the proximity between a locus ***x*** and a gene, under the Weibull distribution assumption (Supplementary Fig. 2).

Epigenomics and in particular, methylation profiles have been integrated to explain tissue specific regulatory mechanisms(15). Weibull distribution for modeling how the minimal distance to a methylated region found in RME database is compared to distance to all the other methylated regions (Supplementary Fig. 3). If it happens that the considered position ***x*** belongs to a methylated region characterized in RME project, we also get the average methylation level for *T* and all the other tissues.

Compared to other approaches mostly relying on RME and/or GTEx, we also considered tissue-specific data sets made available by the research projects focusing on a single tissue. For the brain model, we integrate developmentally differentially methylated positions (dDMPs) (16) found in fetal brain. For the heart model, Heart Enhancer Compendium database(17) for heart development candidates was used.

### Supervised machine learning model training

Considering the aforementioned training sets, we fit several machine learning approaches and compare them based on their 10-folds cross-validated performances (here, area under the ROC curve, AUC), and selected random forest(18) algorithm to train the final model (Supplementary Table 2, AUC = 0.8). Using cross-validation, we optimize both the number of trees and the number of variables to consider at each node in the *randomForest* R package.

### From class probability to rescaled odd-ratio

Current approaches often consider the raw class probability as their functional score, requiring additional tuning step from the user. Here, we propose to rescale the classifier output, into a ready-to-use score. First, we define an optimal cutoff value on the probability (Supplementary Fig. 4a and 5a), as the smallest value which reaches a false discovery rate of 10%. For instance, this threshold is equal to 0.48 for the brain model and to 0.67 for the heart model. Then, we rescale the filtered probability to a score between 0 and 1, using the formula:

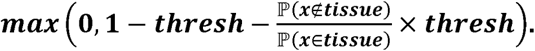

The main advantage of this step is to standardize predictive models, and push loci not tissue-related to a score strictly equal to 0 (Supplementary Fig. 4b and 5b).

### Reference genome

Analyses performed in this study used the hg19 reference genome.

### Evaluation framework

In all analyses presented in this study, we carefully removed positions that were both found in the TiSAn training and the validation sets, avoiding over-optimistic performances.

### GWAS prioritization in coronary artery disease (CAD) cohort

CARDIoGRAM consortium GWAS meta-analysis summary statistics for 8,443,810 SNPs were downloaded at http://www.cardiogramplusc4d.org/media/cardiogramplusc4d-consortium/data- downloads/cad.additive.Oct2015.pub.zip. Correlation between functional score and association strength (Fig. 3) was obtained by binning in 100 percentile bins on reported association P-value. Then, relative score enrichment is computed for top1% variants and iteratively, until merging all the data. We derived confidence interval for both TiSAn and GenoSkyline by random permutations on the GWAS p-values. On the purpose of ranking variants, we filtered variants not predicted as functional by either TiSAn (zero score), or by GenoSkyline (score < 0.15).

### Variant enrichment in vicinity of ASD genes

Variants found in 960 Simons Simplex Collection (SSC) individuals, including probands and parents were filtered based on their pathogenicity using CADD score. We estimated two different threshold values for coding (>15) and non-coding (>10.7) variants. Those values correspond to the top 10%-ile found in the 1000 Genomes data. We also focused the analysis on variants found in a +/-50,000bp windows around well-supported ASD genes, with more than 20 citations in the June 2016 SFARI gene list at http://gene.sfari.org/autdb/HG_Home.do (Supplementary Table 3). The same filters were applied to variants found in 1000 Genomes (1KG) European ancestry population (Phase 3). Coding and non-coding variants were separated based on their RefSeq(19) function annotation. The relative gain in average score (Fig. 2a and b) is calculated by doing the difference between average score in SSC and in 1KG, divided by the score in 1KG. Cumulative score enrichment for SSC over 1KG variants (Fig. 2c) is obtained by binning all variants from the two datasets based on their score, in 5%-ile groups. Then, average score ratio between the two groups is computed in each bin, and summed in a cumulative way, from the top 5% to all the data.

### Transcription factor binding site (TFBS) enrichment in tissue and cell type

ENCODE project provides a large repository for TFBS location in various cell type contexts. Here, we put together two databases, both available as UCSC Genome Browser tracks, *factorbookMotif*, which contains the location of more than 2 million TFBS across the genome, and *EncodeRegTfbsClustered*, which provides information regarding the cell types where TFBS were observed. Overlapping the two databases results in 1,514,086 unique TFBS found in 53 families. For each of those TFBS, we expanded their location using a 1,000 base pairs window, and TiSAn heart and brain scores were extracted. Scores were centered and scaled around the center value and show the actual score enrichment along the window. An average profile was computed for all TFS categories and cell types.

### Software availability

- Genome-wide TiSAn score databases are available in bed format (with index) at:

- http://flamingo.psychiatry.uiowa.edu/TiSAn/TiSAn_Brain.bed.gz
- http://flamingo.psychiatry.uiowa.edu/TiSAn/TiSAn_Heart.bed.gz
- http://flamingo.psychiatry.uiowa.edu/TiSAn/TiSAn_Brain.bed.gz.tbi
- http://flamingo.psychiatry.uiowa.edu/TiSAn/TiSAn_Heart.bed.gz.tbi
- Tutorial and vignettes are also available at http://github.com/kevinVervier/TiSAn.

- GenoSkyline approach: we downloaded brain and heart models on the tool website (http://genocanyon.med.yale.edu/GenoSkyline), in November 2016.

- Combined Annotation Dependent Depletion (CADD): Deleteriousness annotation were performed using the CADD v1.0 (published version) at http://krishna.gs.washington.edu/download/CADD/v1.0/whole_genome_SNVs.tsv.gz

## Results

### Machine learning for predicting tissue-specific functional annotation

The design of TiSAn models is outlined in Figure 1 (details in Materials & Methods section). Taking advantage of publically available datasets(5,6), we extracted more than 350 different genome-wide variables which were used to describe two large sets of disease-related loci. Training a supervised machine learning model requires positive and negative examples: here, positive examples were nucleotide positions that had been previously linked to a tissue-specific disease, and negative examples were variants that had no established link to the tissue-specific disease in question. Predictive models were trained on the labeled datasets and optimized to achieve high discrimination of tissue-specific loci (Sup. Fig. 4-5). Here, a position with a score equal to 1 can be considered strongly associated with the tissue, whereas a score of 0 means no association at all, and such a position is usually discarded in subsequent analysis.

**Figure 1:**
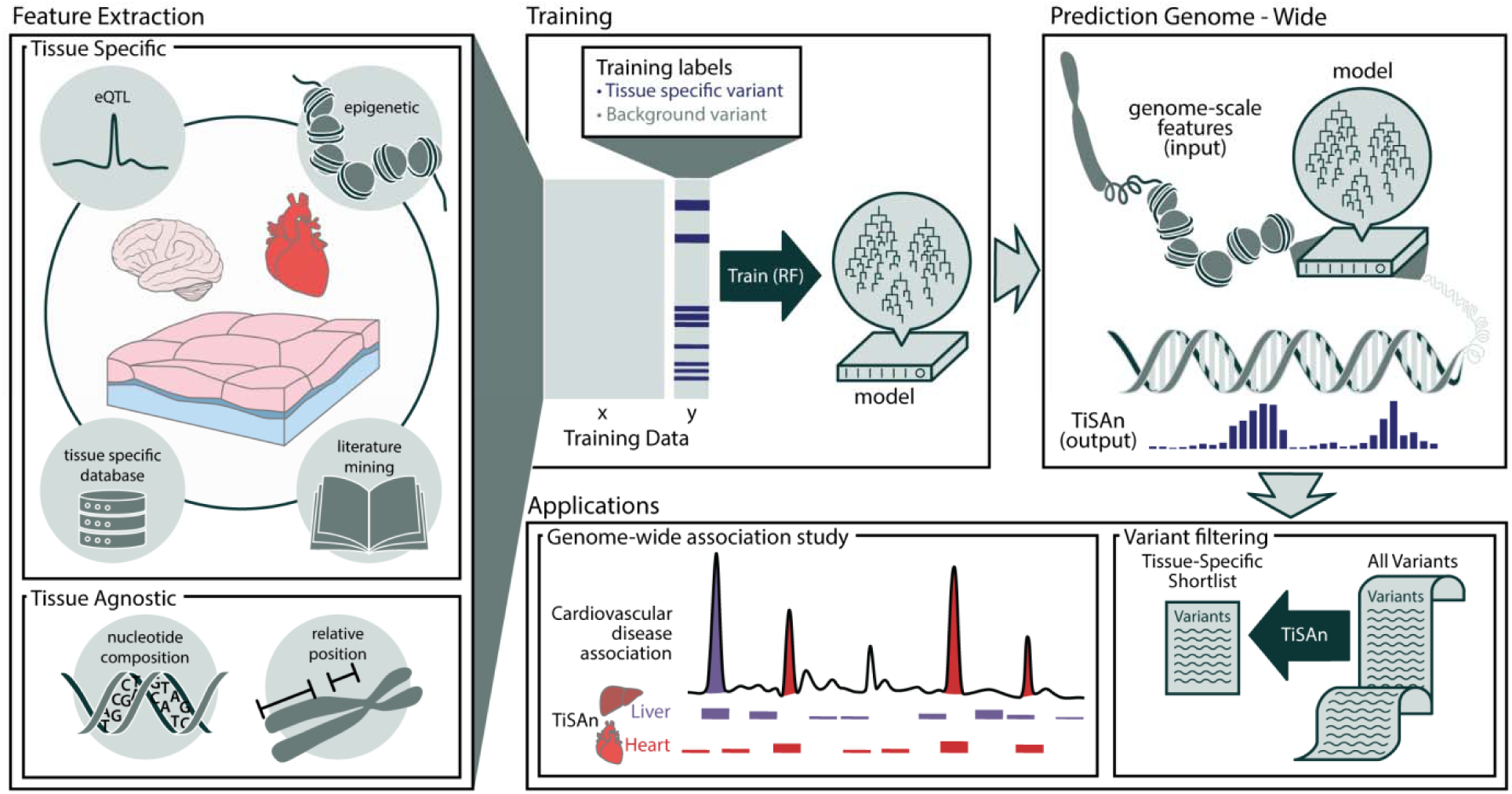
TiSAn framework overview. *Each nucleotide position in the genome is annotated with multiple levels of omics information, such as sequence content, methylation level, proximity to genes, etc. (see Methods). This information is extracted for training sets, comprised of deleterious variants with or without an association with the tissue of interest. Using supervised machine learning, specifically a Random Forest (RF), a predictive model combines each feature with respect to its ability to predict whether a position will be functionally associated with the tissue of interest. Model output consists in a tissue-specific functional score spanning between no functional relevance to the tissue (0) and strong functional relevance to the tissue (1). This score can then be used, for instance, to filter down large lists of candidate variants for further investigation, or to isolate the contribution of different tissues to a complex trait.*

In the following, we demonstrate TiSAn performance in three different settings, i) performing tissue-specific enrichment in case-control cohort, ii) enhancing results from a genome-wide association study (GWAS), iii) extracting genome-wide tissue-specific transcription factors. We also consider a recently proposed tool, GenoSkyline(7) that provides a genome partition in terms of functional segments, using methylation data only. Our approach aims to provide functional prediction at the single nucleotide resolution, because variants found in large predicted functional blocks (as is the case in GenoSkyline) may in fact have different functional effects.

### Brain-specific variant prioritization in a sample with familial risk for autism

#### Genome-wide enrichment for brain-related variations in affected individuals

The Simons Simplex Collection (SSC, http://base.sfari.org) provides whole genome sequencing for one of the largest autism spectrum disorder (ASD) cohorts currently available. We hypothesized that deleterious genetic variation found in the vicinity of ASD-related genes would show higher enrichment in terms of brain-related functional consequences (as measured by the TiSAn-brain and GenoSkyline-brain scores) in the SSC compared to the 1000 Genomes (1KG)(20). We further assessed enrichment using the respective heart-specific scores as a form of negative control. In this analysis, the TiSAn-brain score shows the only positive tissue-specific enrichment, over 50% for coding variants (Fig. 2A) and around 10% for non-coding variants (Fig. 2B). Notably, there is a significant difference between TiSAn brain and heart scores (Wilcoxon signed-rank test, *P < 2x10*^*-16*^), suggesting effective tissue specificity, whereas this was not observed for GenoSkyline models (Wilcoxon signed-rank test, *P = 0.351*).

**Figure 2:**
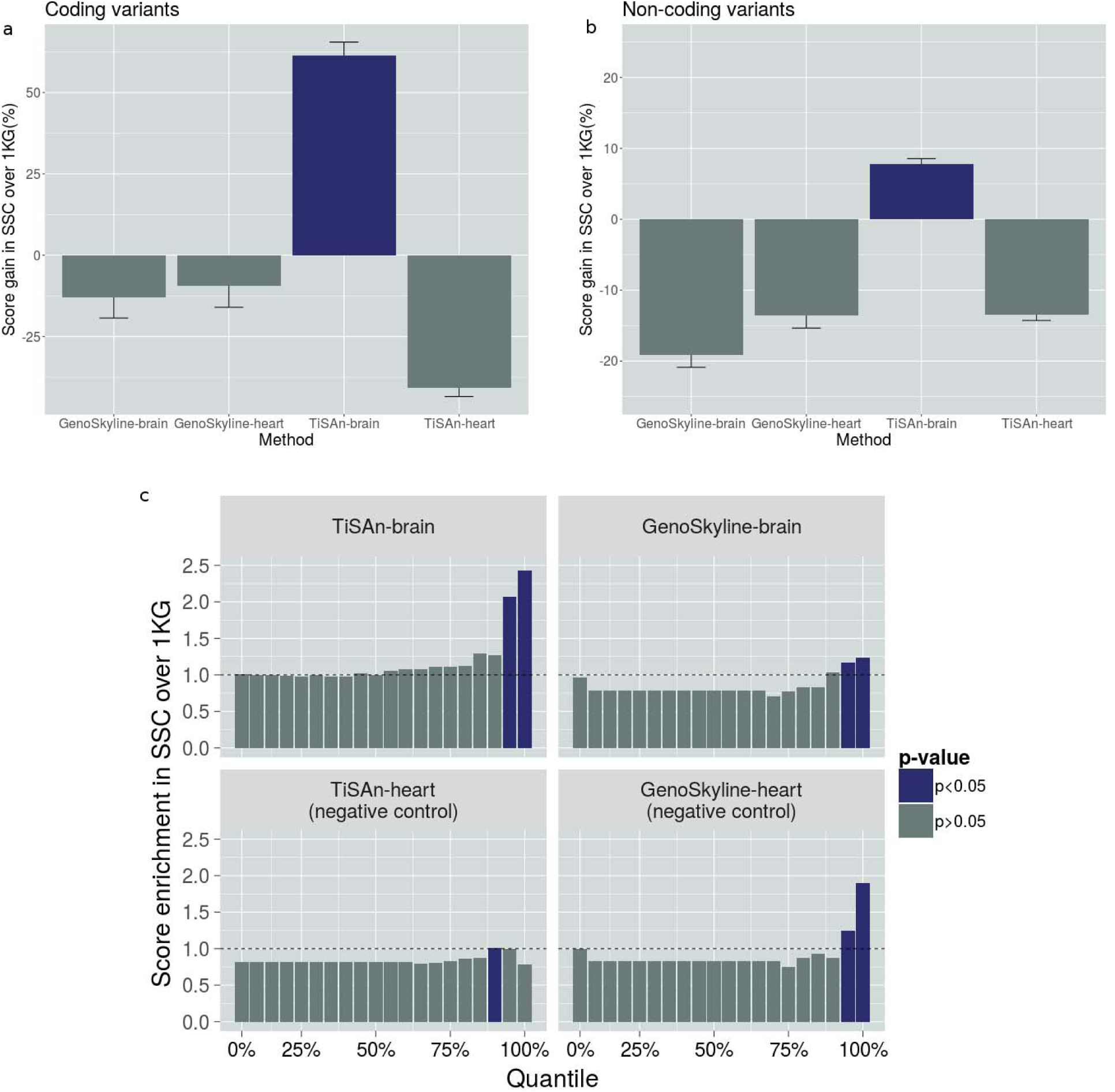
Brain-related functional enrichment in a case-control setting. *Comparison of Simons Simplex Collection (SSC) variants with 1000Genomes (1KG) variants. Coding variants (a) and non-coding variants **(b)**. Both brain and heart models for TiSAn and GenoSkyline were evaluated. **(c) Functional score enrichment in SSC variants compared to 1KG variants.** After sorting SSC and 1KG variants based on their score, we compute cumulative enrichment for each 5%-ile. Blue bars correspond to significant difference between SSC and 1KG, using the Chi-squared test (p-value <0.05).*

#### Case-control variants filtering with brain-specific annotation

Next, we ranked and binned variants according to their tissue-specific scores (i.e., TiSAn or GenoSkyline) and calculated the enrichment of SSC deleterious variants in each bin, with respect to deleterious 1KG variants. Because the SSC is a neurodevelopmental cohort, we expect to see over-representation of SSC variants in the most confidently called brain-related genomic regions. Indeed, significant enrichment of SSC variants was observed in the top quantiles for TiSAn-brain but also for both GenoSkyline models (Fig. 2C). Surprisingly, the GenoSkyline-heart model reports a more pronounced enrichment than the corresponding brain model, suggesting a potential lack of tissue specificity for GenoSkyline. TiSAn-brain achieves the highest enrichment by ranking 2.5 times more SSC variants in the top 5% than 1KG variants.

#### Autism and calcium channel genes

An autism-related calcium voltage-gated channel gene, *CACNA1C*(21), was the gene with the highest TiSAn-brain score enrichment in SSC data, suggesting that deleterious variants at this locus are more likely to affect brain function. In particular, we identified 116 non-coding deleterious variants (see methods for CADD thresholds) in CTCF(22) transcription factor binding sites around *CACNA1C* location in SSC data, while none were found in unaffected population. Mutations in the same region also hit non-coding RNAs (ncRNAs) more frequently in SSC population than in control population (Fisher’s exact test, *P < 2x10*^-16^). Interestingly, 5 of these ncRNAs (Supplementary Table 4) were found in linkage disequilibrium with loci associated with autism and Tourette’s syndrome according to the LincSNP database(9).

### Heart-related signal prioritization in coronary artery disease

#### Genome-wide association strength and annotation score

Current approaches to GWAS analysis rely mostly on association strength (e.g., *P*-value) to prioritize candidate regions. These variants often belong to large linkage-disequilibrium (LD) blocks, making it difficult to decipher the actual causal genetic mechanism. Here, we apply TiSAn to the Coronary Artery Disease (CAD) CARDIoGRAM consortium GWAS meta-analysis(23), and we demonstrate that the TiSAn-heart score is significantly higher among the most associated variants (Fig. 3, (Student t-test, *P < 2x10*^-16^)). Furthermore, the top 100 SNPs (according to their *P*-value) with a non-zero TiSAn were all found in LD with genomic regions strongly associated with coronary artery disease, demonstrating TiSAn high sensitivity. In this analysis, no significant enrichment was observed for GenoSkyline-heart (Wilcoxon signed-rank test, *P = 0.12*) or brain models (Supplementary Fig. 8).

**Figure 3:**
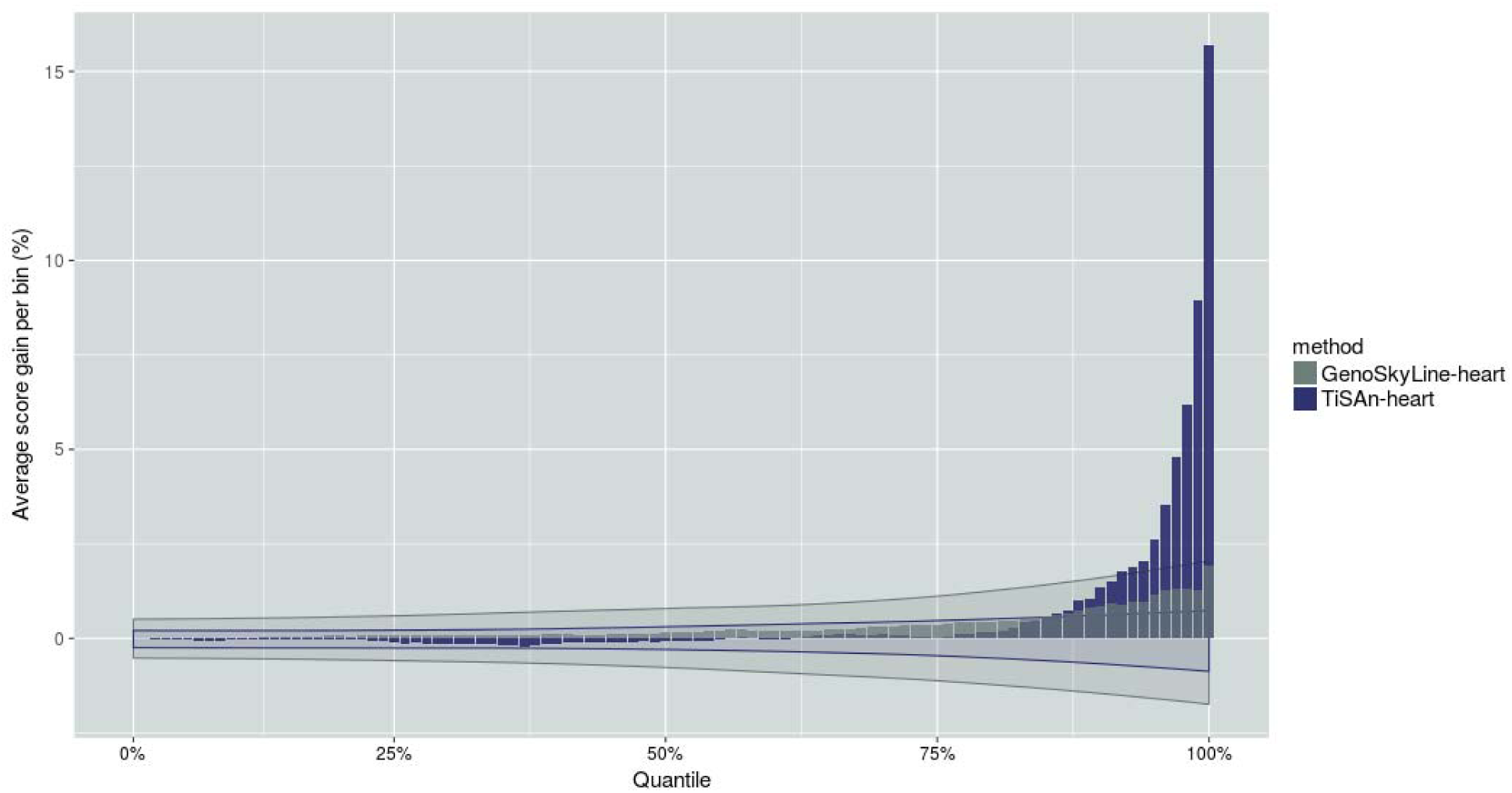
CAD-GWAS signal prioritization using heart-related models. *Genetic variants were binned by percentiles, based on their association P-values. In each of those bins, we report average functional scores (blue: TiSAn-heart, grey: GenoSkyline-heart). Shaded areas represent confidence interval for the corresponding method, after GWAS P-value random permutations.*

#### Reduction of multiple hypothesis burden in GWAS

We filtered tissue-relevant genotyped variants before the GWAS analysis, so that only heart-related variants would be considered in the correction for multiple testing. In the case of CAD-GWAS, this reduced the number of SNPs considered by 75%. This resulted in significant loci being narrowed by 20% on average (paired Student t-test, *P = 0.019*). Furthermore, the overall enrichment in transcription factor binding sites (TFBS) among significant loci is conserved between the original set of and the TiSAn-filtered one (Chi-squared test, *P = 0.51*), suggesting that the regulatory content is preserved after the filtering step (a further analysis of TFBS, provided in the supplementary information, suggests that TiSAn can reveal which TFs have important functional roles in specific tissues). Reducing the number of tested variants directly recalibrates the multiple-testing correction threshold used to determine significant loci from 5x10^−8^ to 1.6x10^−7^. Here, 91 new loci were found significantly associated with coronary artery disease, and show a significant enrichment in EBF1 TFBS (Fisher’s Exact test, P = 3.2x10^−6^), which has been previously linked to obesity, diabetes, and cardiovascular disease(24).

## Discussion

Integrative approaches like TiSAn hold great promise for helping genomics researchers narrow down massive lists of variants to focus on those that are most relevant to the tissue or disease at hand. However, few such tools currently exist, with most development efforts focusing on improving estimators of general (and not tissue-specific) deleteriousness(25). GenoSkyLine, a recently developed tool that utilizes genome-scale tissue-specific epigenetic data, allowed us to benchmark TiSAn and demonstrate its effectiveness in prioritizing genetic variants that are most likely to play a role in the tissue-specific disease processes under consideration.

Specifically, we showed that individuals with elevated risk for autism (i.e., probands and their family members) had more deleterious WGS variants that were predicted to be brain relevant (by TiSAn-Brain) than controls. At the same time, we were unable to show any such differences in regions identified as brain-relevant by GenoSkyLine. Specificity was demonstrated in this analysis by the absence of a similar case/control difference in deleterious variants predicted (by TiSAn-Heart) to be relevant to cardiovascular tissue. Further, we showed that strongly associated GWAS hits in a study of coronary artery disease have a significantly higher TiSAn-Heart signal than non-associated SNPs, supporting our method’s ability to correctly prioritize tissue-specific variants. Again, we were unable to observe this difference using the GenoSkyLine score for cardiovascular tissue. We also demonstrated the practical advantages of reducing GWAS multiple testing burden by pre-filtering SNPs on the basis of their estimated tissue relevance. In each of these analyses, TiSAn showed an ability to correctly prioritize variants according to tissue specific action, while we were unable to do so with GenoSkyLine, the current state of the art for this application. TiSAn thus represents an important development towards leveraging the massive amount of underutilized information (i.e., non-coding variation) coming from whole genome sequencing studies.

Several technical points related to the development of TiSAn are worth mentioning. First, a comparison between multiple machine learning algorithms (Supplementary Table 2) led us to use random forests, known to better handle non-linearity and correlation between variables. Recently, deep learning has been evaluated in the context of variation effects on chromatin(11), and future analyses will investigate the impact of using this algorithmic framework. Another issue is that supervised learning requires genomic positions with accurate class labels, in this case, known to be either associated with a given tissue or not. However, for most of the available data, such a label does not exist, especially for positive association with a tissue-specific trait. Imbalance-aware machine learning(26) could be a solution to efficiently train predictive models in the case of underrepresented classes.

Researchers interested in other tissues beyond brain or heart can derive their own functional annotation for a selected tissue of interest (Online Methods), and we have provided thorough documentation, including tutorials, on how to use TiSAn in typical genome informatics workflows.

## Acknowledgments

Data on autism spectrum disorder variation have been contributed by Simons Simplex Collection investigators and have been downloaded from http://sfari.org/resources/sfari-base.

Data on coronary artery disease have been contributed by CARDIoGRAMplusC4D investigators and have been downloaded from www.cardiogramplusc4d.org.

## Supplementary Materials

### Tissue-specific signal in transcription factor binding sites

#### Genome-wide enrichment in transcription factor binding sites

Transcription factor (TF) binding sites (TFBS) are associated with observed differences in gene expression across tissues(12). We hypothesized that computing TiSAn score profiles in TFBS could provide insight about the tissue-related action of specific TFs. The ENCODE project provides TFBS detection in 80 different cell types for more than 50 TFs. TiSAn scores were predicted for millions of loci using a 1,000bp window centered on TFBS. Average TiSAn profiles for each TF allow us to identify the sites showing an overall enrichment across cell types. For instance, strong heart-related signal was found among TFBS for BHLHE40, CEBPB, FOXA1, GATA1, HNF4A, JUN, MAFK, MAX, MYC, POU2F2, STAT1, and TAL1. Notably, CEBPB TFBS are enriched for TiSAn-heart score in 6 cell types, including two related to smooth muscles (A549 and IMR90) and one related to liver (HepG2) (Supplementary Fig. 7A), and *CEBPB* has been associated with cardiac hypertrophy(27) and fatty liver disease(28).

#### Brain-specific regulatory features

Brain-specific binding patterns in mouse have been observed for CTCF, the major regulator of chromatin state(22), and our analysis demonstrates a consistent signature with TiSAn-Brain enrichment at the CTCF binding locations across 70 different cell types (Supplementary Fig. 8), suggesting the importance of chromatin conformation in brain development and function. Functional enrichment patterns were also found for the critical brain transcription factor REST in 10 different cell types (Supplementary Fig. 7B). Among these were 3 brain cancer cell lines (U87, SK-N-SH, and PFSK-1), which support recent findings on the importance of REST in neuroblastoma drug sensitivity(29).

**Supplementary Figure 1:**
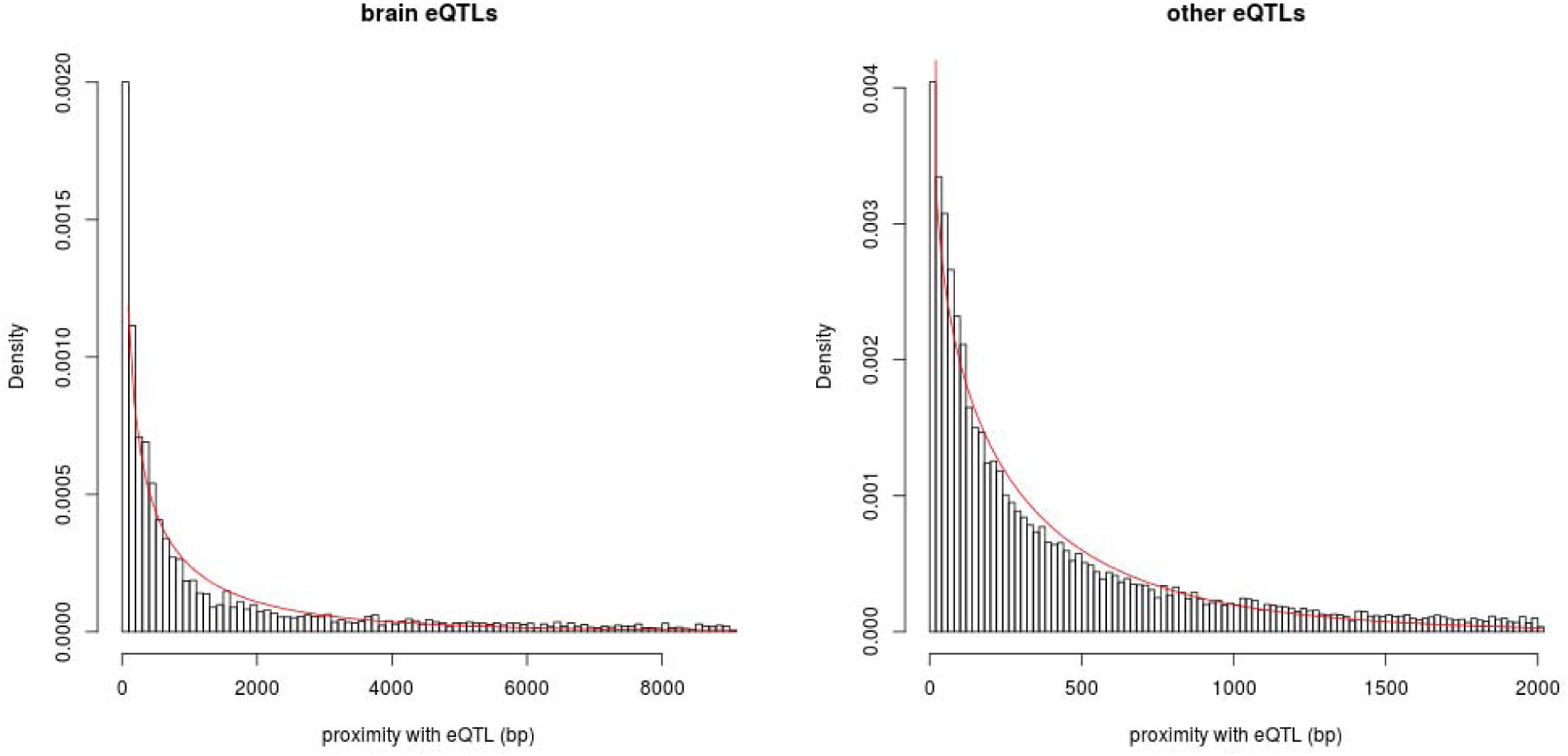
*Distribution of distance to the closest GTEx expression quantitative trait locus (eQTL) for brain (left) and non-brain (right) tissues. The red lines correspond to a Weibull distribution fit. Estimated parameters for left (resp. right) figure are: shape = 0.351 (resp. 0.315) and scale = 21,888 (resp. 9,111).*

**Supplementary Figure 2:**
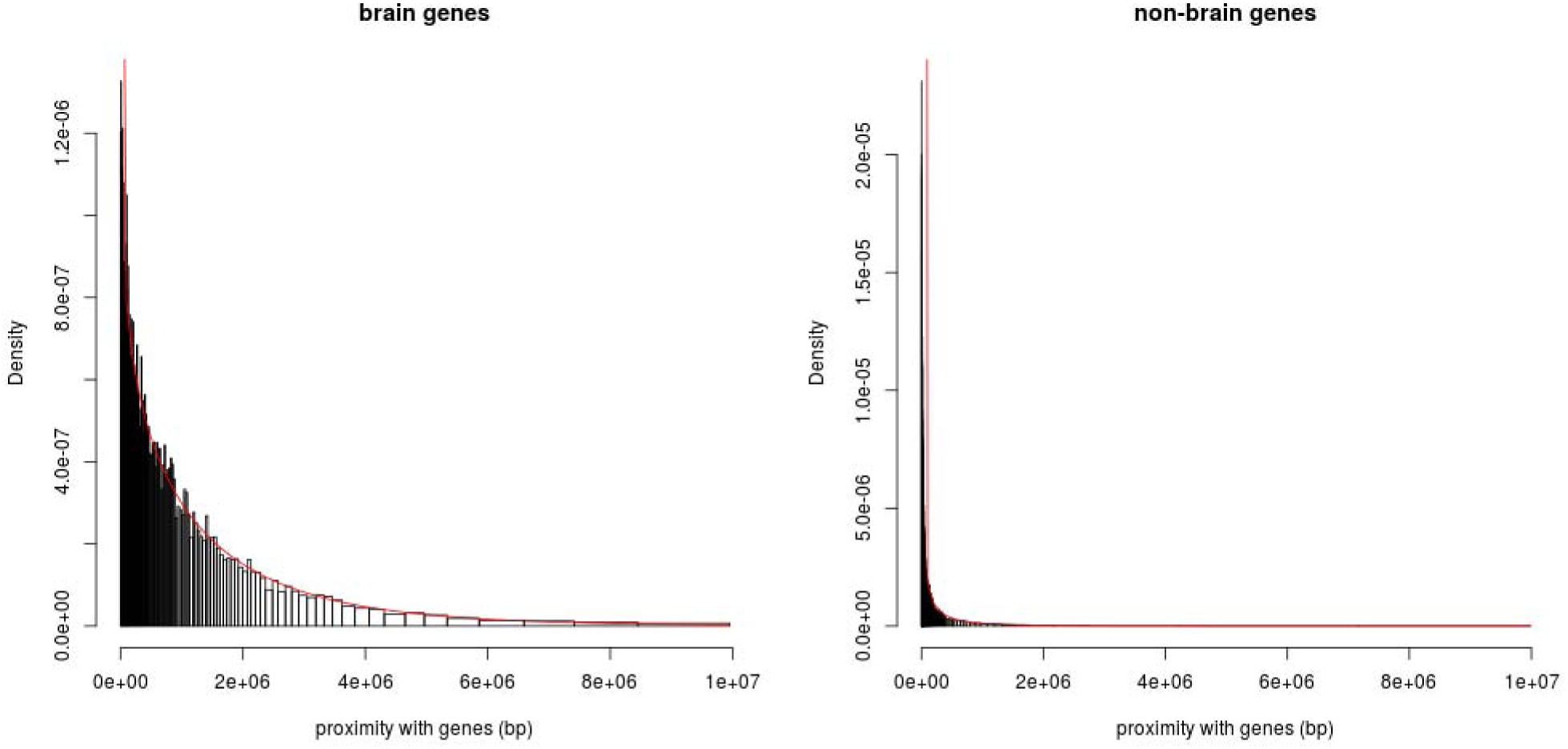
*Distribution of distance to the closest gene for brain (left) and non-brain (right) tissues. The red lines correspond to a Weibull distribution fit. Estimated parameters for left (resp. right) figure are: shape = 0.852 (resp. 0.529) and scale = 1,453,217 (resp. 201,985).*

**Supplementary Figure 3:**
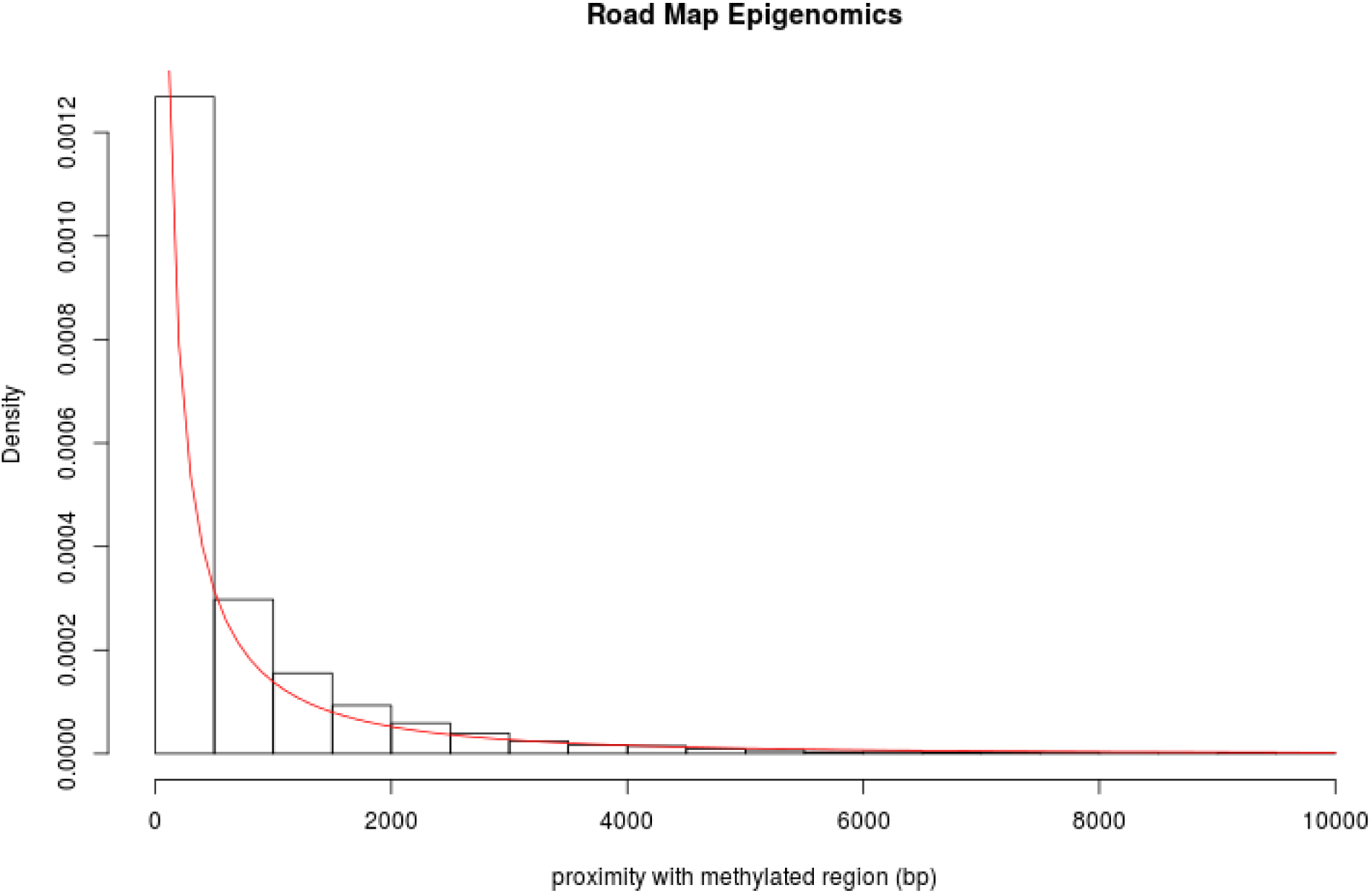
*Distribution of distance to the closest methylated region found in RoadMap Epigenomics database. The red line corresponds to a Weibull distribution fit. Estimated parameters are: shape = 0.746 and scale = 590.3.*

**Supplementary Figure 4:**
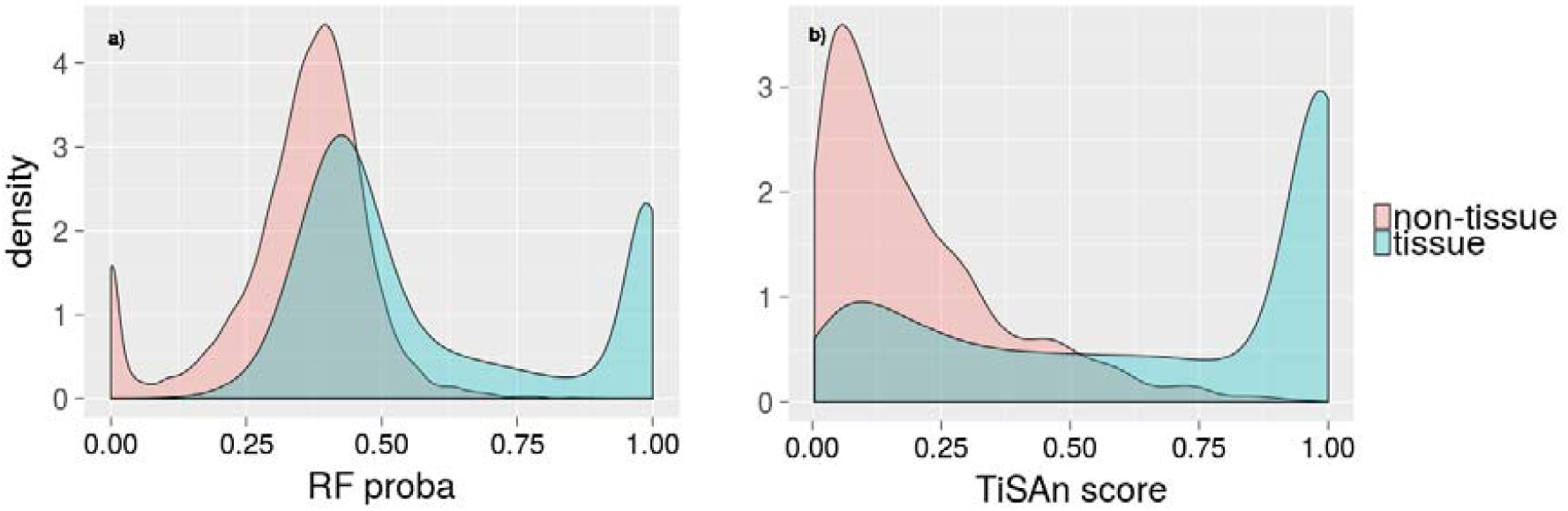
TiSAn-brain. *cross-validation performances. (a) Random forest raw output distribution. (b) TiSAn score obtained after rescaling odd-ratios.*

**Supplementary Figure 5:**
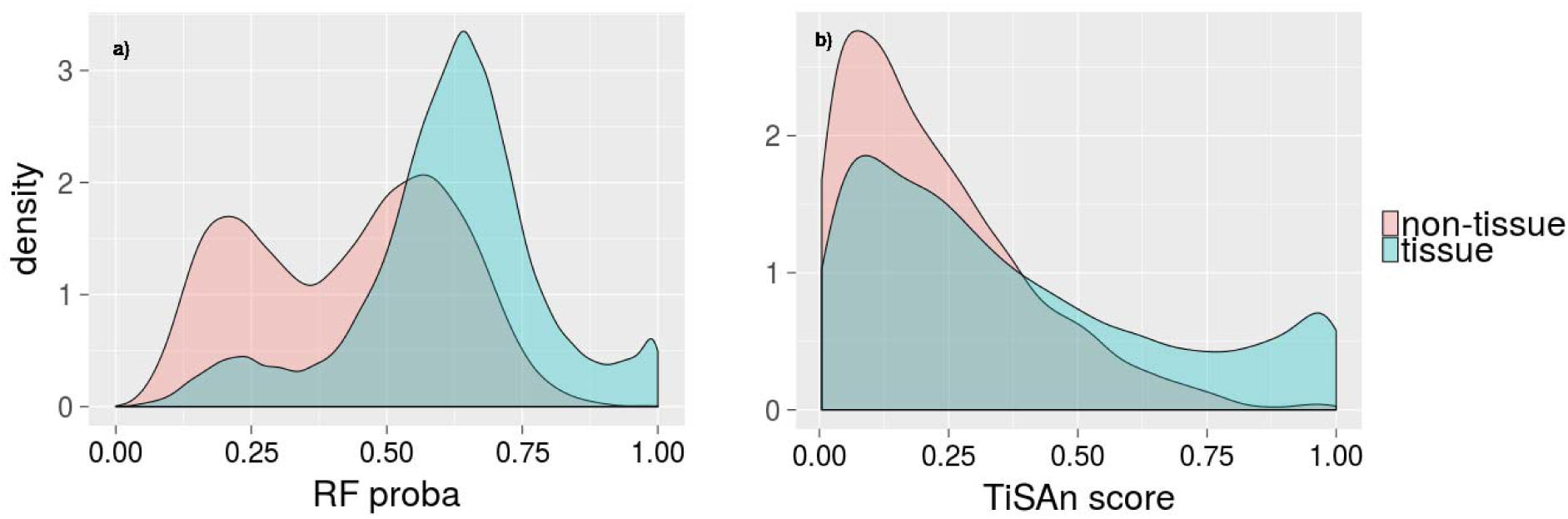
TiSAn-heart. *cross-validation performances. (a) Random forest raw output distribution. (b) TiSAn score obtained after rescaling odd-ratios.*

**Supplementary Figure 6:**
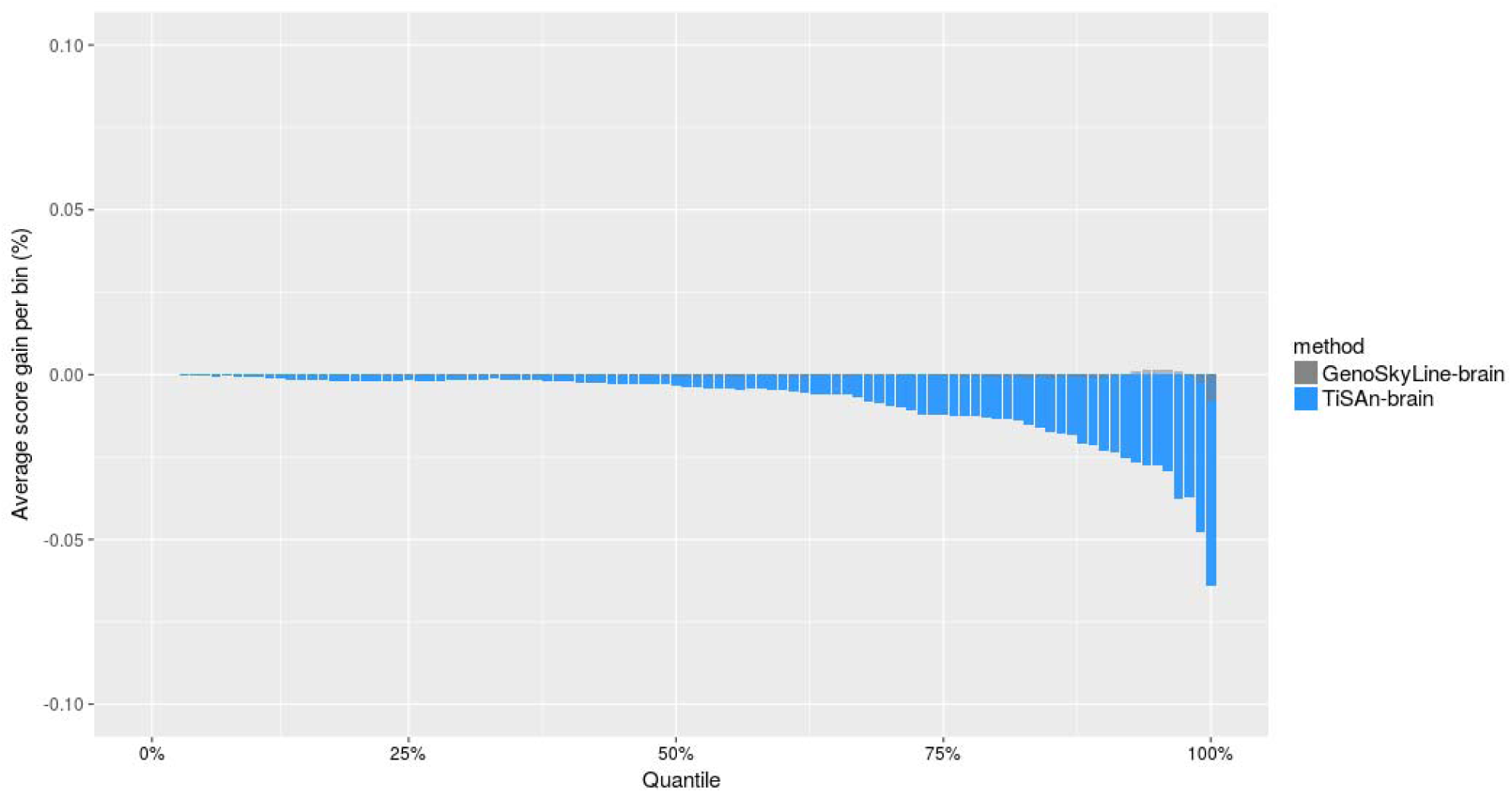
*CAD-GWAS signal prioritization using brain-related models. Genetic variants were binned by percentiles, based on their association p-values. In each of those bins, we reported average functional scores (blue: TiSAn-brain, grey: GenoSkyline-brain)*

**Supplementary Figure 7:**
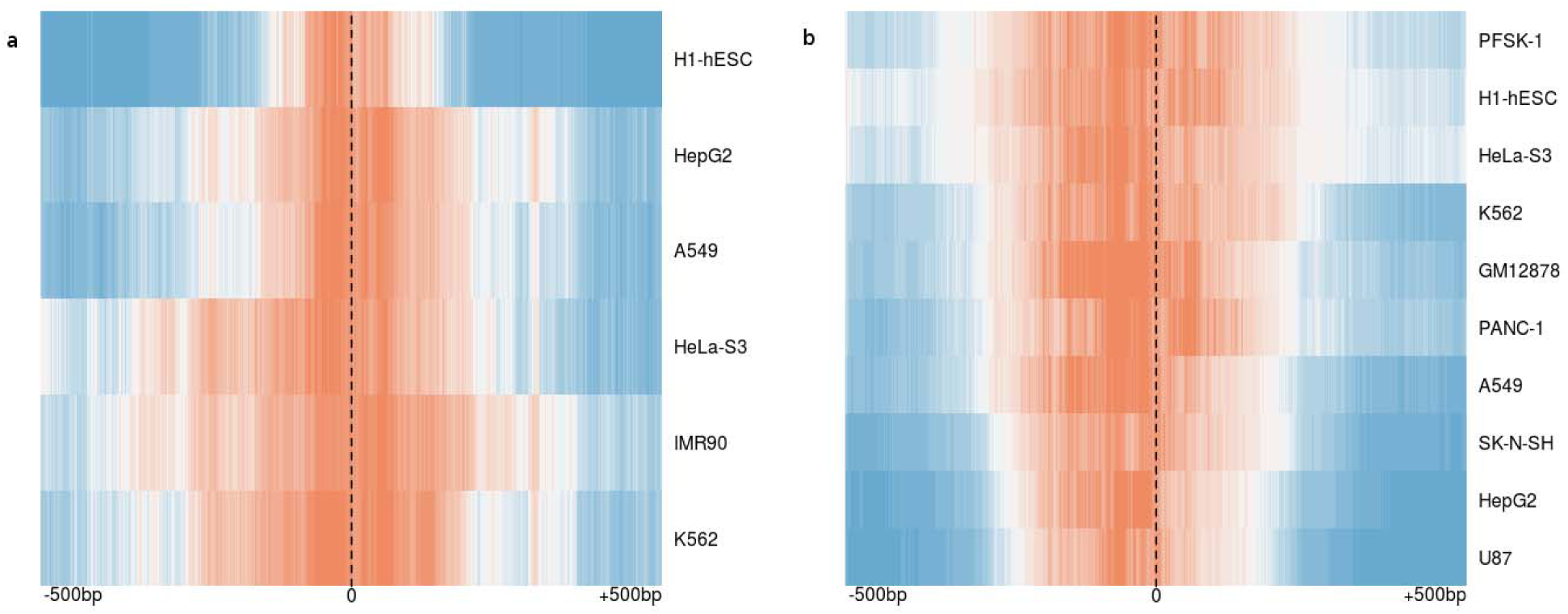
Genome-wide tissue-specific transcription factor binding sites (TFBS) characterization. *Functional score profiles were obtained using a 1,000bp window centered on the TFBS (dash line). Positive enrichment (orange) and negative enrichment (blue) are reported for each different cell type in row. **(A) TiSAn-heart enrichment in CEBPB TFBS**. Locations for ENCODE TFBS were found in 6 different cell types. **(B) TiSAn-brain enrichment in REST TFBS.** Locations for ENCODE TFBS were found in 10 different cell types.*

**Supplementary Figure 8:**
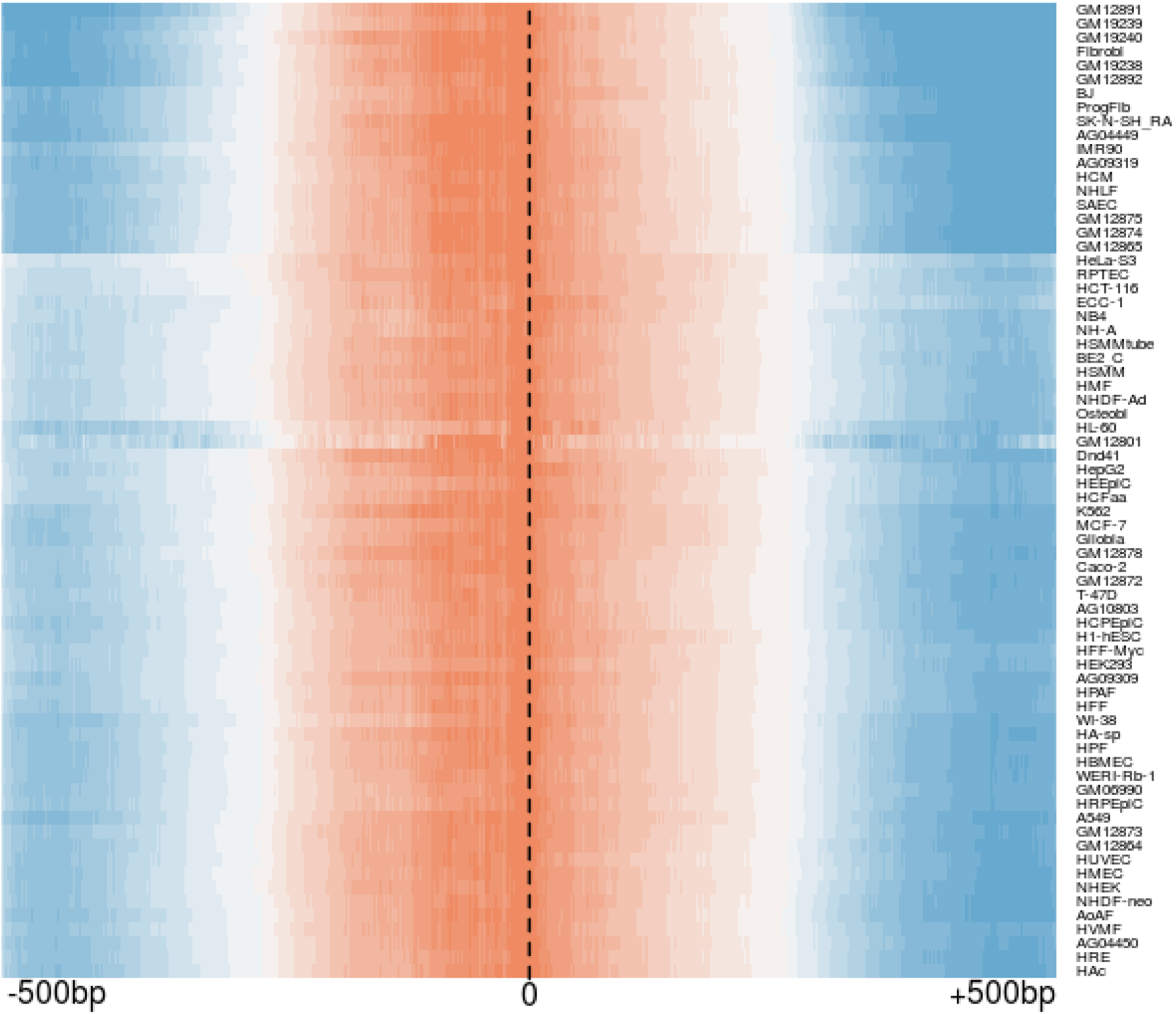
*TiSAn-brain enrichment in CTCF transcription factor binding sites (TFBS). Locations for ENCODE TFBS were found in 70 different cell types. Functional score profiles were obtained using a 1,000bp window centered on the TFBS (dash line). Positive enrichment (orange) and negative enrichment (blue) are reported for each different cell type in row.*

**Supplementary Table 1:** Training set composition. *For both heart and brain tissues, we report the count of positive and negative examples used to train TiSAn models. The counts are divided in two parts, corresponding to variants found in large intergenic non-coding RNAs database, or in* genotype array probesets.

| Tissue | Category | Count |
| --- | --- | --- |
| Brain | Positive | 10,715 (5,535 + 5,180) |
| Brain | Negative | 22,811 (13,861 + 8,950) |
| Heart | Positive | 28,473 (9,760 + 18,713) |
| Heart | Negative | 62,438 (22,476 + 39,962) |

**Supplementary Table 2:**
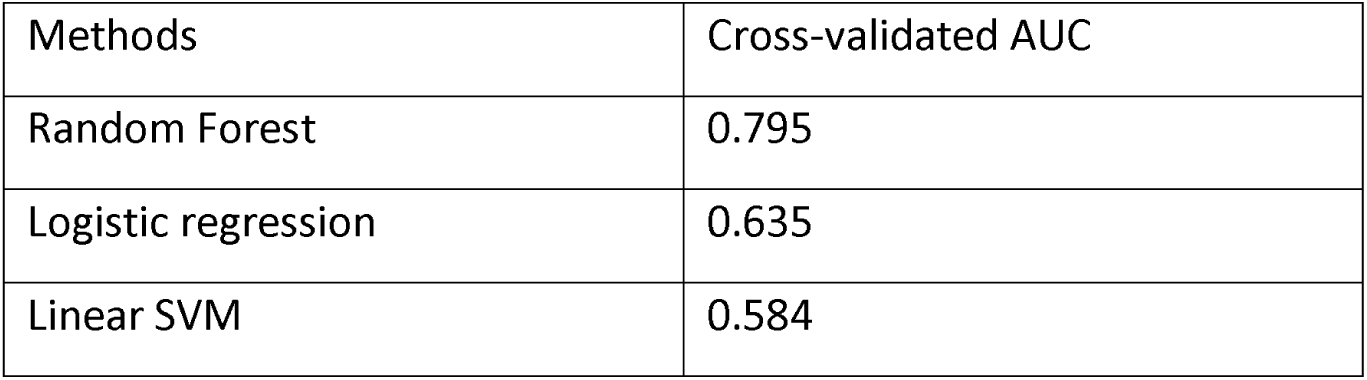
*10-folds cross-validated performances obtained during TiSAn-brain model training, for different classification strategies. AUC: Area under ROC curve.*

**Supplementary Table 3:** *List of SFARI autism-related genes, supported by literature.*

| Gene | Citations | Gene | Citations |
| --- | --- | --- | --- |
| *NRXN1* | 61 | *GABRB3* | 30 |
| *CNTNAP2* | 49 | *SCN2A* | 30 |
| *SHANK3* | 49 | *FOXP2* | 28 |
| *PTEN* | 39 | *RBFOX1* | 28 |
| *CACNA1C* | 35 | *SYNGAP1* | 28 |
| *OXTR* | 34 | *AUTS2* | 27 |
| *RELN* | 34 | *GRIN2B* | 25 |
| *MET* | 32 | *DPP6* | 22 |
| *DISC1* | 31 | *MBD5* | 22 |
| *SCN1A* | 31 | *SLC6A4* | 22 |

**Supplementary Table 4:** *Non-coding RNAs found in linkage disequilibrium with neurodevelopmental and psychiatric disorders*

| **LincSNP ID** | **Chr** | **Start** | **End** | **Associated disorder** |
| --- | --- | --- | --- | --- |
| *LSLNC096364* | 12 | 2907933 | 2909631 | *Suicide attempts in bipolar disorder* |
| *LSLNC141017* | 12 | 2901746 | 2904044 | *Autism with low IQ* |
| *LSLNC169768* | 12 | 2901745 | 2904044 | *Autism with low IQ* |
| *LSLNC169511* | 12 | 1936729 | 1940267 | *Tourette's syndrome* |
| *LSLNC027117* | *12* | *1936728* | *1940267* | *Tourette's syndrome* |

